# Visual dysfunction predicts cognitive impairment and white matter degeneration in Parkinson’s disease

**DOI:** 10.1101/2020.10.12.335547

**Authors:** Angeliki Zarkali, Peter McColgan, Louise-Ann Leyland, Andrew J. Lees, Rimona S. Weil

## Abstract

Visual dysfunction predicts dementia in Parkinsons disease (PD), but whether this translates to structural change is not known. We aimed to identify longitudinal white matter changes in patients with Parkinsons disease and low visual function and also in those who developed mild cognitive impairment (MCI). We used fixel-based analysis to examine longitudinal white matter change in PD. Diffusion MRI and clinical assessments were performed in 77 patients at baseline (22 low visual function /55 intact vision; and 13 MCI, 13 MCI converters /51 normal cognition) and 25 controls and again after 18 months. We compared micro-structural changes in fibre density, macro-structural changes in fibre bundle cross-section (FC) and combined fibre density and cross-section across white matter, adjusting for age, gender and intracranial volume. Patients with Parkinsons and visual dysfunction showed worse cognitive performance at follow up and were more likely to develop MCI compared with those with normal vision (p=0.008). Parkinsons with poor visual function showed diffuse micro-structural and macro-structural changes at baseline, whereas those with MCI showed fewer baseline changes. At follow-up, Parkinsons with low visual function showed widespread macrostructural changes, involving the fronto-occipital fasciculi, external capsules, and middle cerebellar peduncles bilaterally. No longitudinal change was seen in baseline MCI or in MCI converters, even when the two groups were combined. Parkinsons patients with poor visual function show increased white matter damage over time, providing further evidence for visual function as a marker of imminent cognitive decline.

## Introduction

The risk of dementia is markedly increased in Parkinson’s disease (PD) but its onset and severity is highly heterogeneous^1,2^, making individual-level predictions difficult and limiting timely therapeutic interventions. There is now a growing body of evidence that Parkinson’s patients with visual dysfunction are at a greater risk of dementia^3–5^, but whether this translates into structural change over time is not yet known.

Animal and cell models suggest that axonal injury is an early event in the degenerative processes leading to Parkinson’s dementia^6–8^. In patients with PD, white matter changes are seen and can be measured using diffusion-weighted imaging. These increase with worsening cognition and may precede grey matter atrophy.^9,10^ However, white matter imaging using conventional diffusion-tensor imaging metrics is insensitive and cannot accurately model crossing fibres, which affects a large proportion of white matter tracts.^11^

Recently, higher-order diffusion models, such as Fixel-based analysis (FBA) have emerged as a more sensitive and fibre-specific framework for identifying white matter alterations.^12^ FBA quantifies changes in specific fibre populations within a voxel known as a “fixel”, thus allowing comparisons of specific tracts rather than metrics averaged across voxels.^12^ We have recently shown that FBA is more sensitive than standard techniques in identifying white matter changes in patients with PD^13^. A recent FBA study showed macro-structural changes within the corpus callosum in patients with Parkinson’s disease which worsened with disease progression. but there were only limited cognitive assessments and it was not designed to examine white matter changes that relate to cognitive progression in PD^14^.

We have examined the white matter changes that evolve during the early stages of cognitive impairment in PD. We assessed patients with PD and visual dysfunction, who are known to be at risk of incipient dementia^1,15,16^ and examined white matter alterations at baseline and after 18 months follow-up, comparing them to PD patients with intact visual function. We also examined white matter changes linked with PD and mild cognitive impairment (PD-MCI) in the same group of patients. We assessed white matter changes in patients with PD-MCI at baseline, and longitudinally in PD-MCI and in patients who progressed to develop PD-MCI. We hypothesised that patients with Parkinson’s and visual dysfunction would show worsening cognitive performance and diffuse white matter degeneration at follow up, compared to those with intact vision; and that more diffuse changes would be evident in this group than in patients with PD-MCI and PD-MCI converters, reflecting the greater sensitivity of poor vision as a predictor of degeneration in PD.

## Methods

An overview of the study methodology is seen in Figure 1.

**Figure 1.**
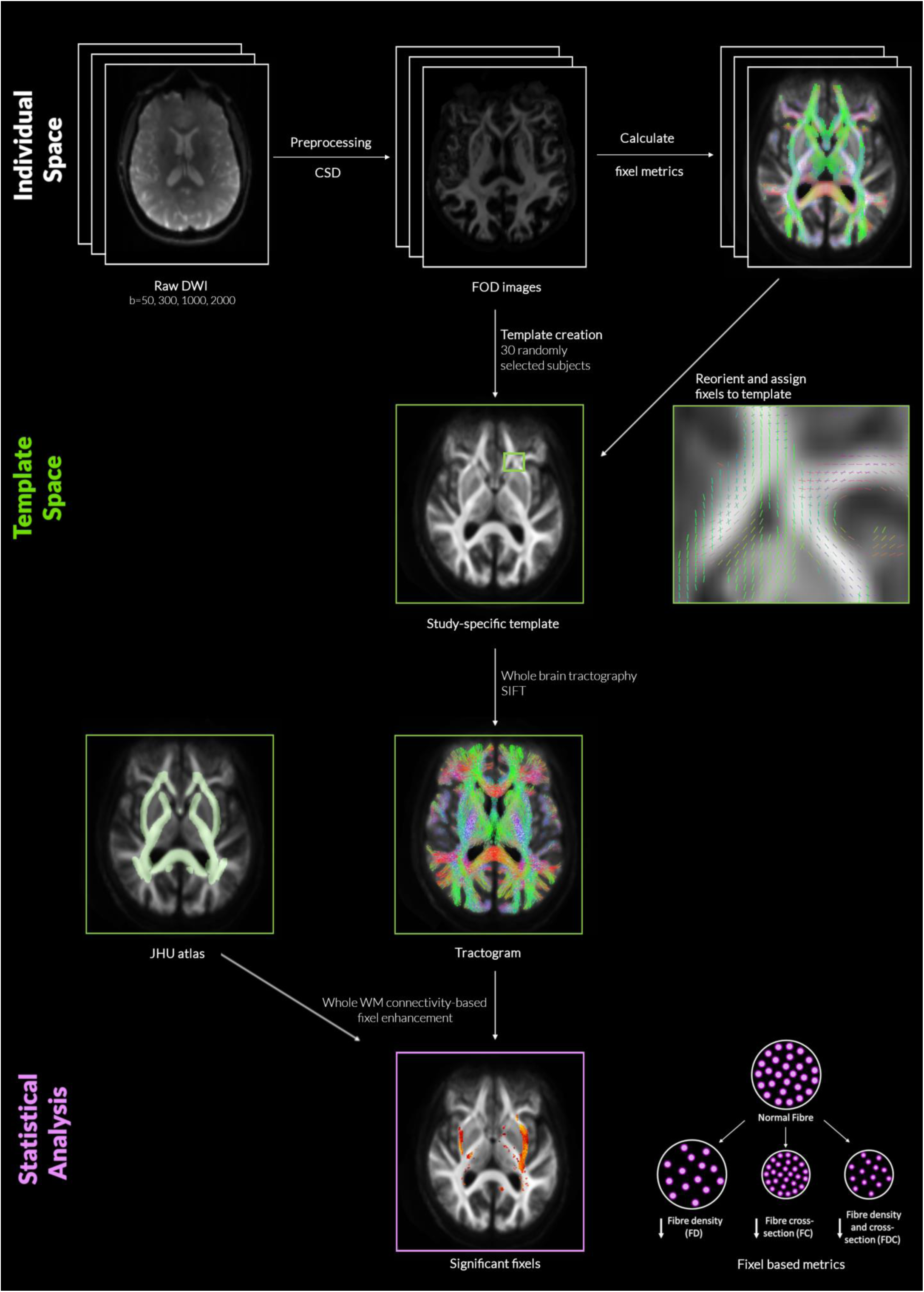
Overview of study methodology (opposite) Individual raw diffusion-weighted imaging scans (DWI) underwent preprocessing and constrained spherical deconvolution (CSD) leading to estimation for distribution of fiber orientation (FOD) and calculation of the three fixel metrics for each participant. A study-specific template was created at baseline from 30 randomly selected participants (20 Parkinson’s disease and 10 controls). Fixels from each subject were then registered to the template fixels. Whole brain probabilistic tractography with 10 million streamlines was performed on the template image and spherical-deconvolution informed filtering of tractograms was then performed resulting to 2 million streamlines. Connectivity-based Fixel Enhancement was performed on the resulting streamlines for statistical comparison with 5000 permutations and family-wise error correction across all white matter fixels using the JHU atlas as a mask. DWI: diffusion-weighted imaging; CSD: multi-shell multi-tissue constrained spherical deconvolution; FOD:fiber orientation distribution; FD: fiber density; FC: fiber-bundle cross-section; FDC: fiber density and cross-section; JHU: Johns Hopkins University.

A total of 102 participants were included in the study: 76 patients with PD and 25 unaffected controls. Participants underwent clinical and psychological assessments as well as brain imaging at baseline and again after 18 months (Visit 2). Participants with PD were further classified according to their performance in two computer-based visual tasks, the Cats and Dogs task and Biological Motion task. Details of stimulus generation and experimental procedures have been previously described^5,16–18^. Performance in these tasks have been shown to correlate with cognitive but not low-level visual performance at baseline as well as worsening cognition at one-year follow up^5,17^. We have also shown that task performance is associated with fibre-specific white matter changes^13^. Similar to previous work, participants who performed worse than the group median performance on both tasks at baseline were classified as PD low visual performers (n=22). All other participants were classified as PD high visual performers (n=54).

All participants underwent a series of clinical and psychological assessments at both study time points. All PD participants had a MoCA score ≥ 26 at baseline, above the cut-off for Parkinson’s dementia of the Movement Disorder Society Task Force Criteria^19^. PD with Mild Cognitive Impairment (PD-MCI) was defined as impaired performance (<1.5 SD of control performance) on at least two domain neuropsychological tests according to MDS criteria^20^. We examined both PD participants with MCI at baseline (n=13, of whom 9 were low visual performers) and those who had normal baseline cognition but developed MCI at Visit 2 (n=13 of whom 4 were low visual performers), defined here as PD-MCI (n=26), in order to capture the group at risk of declining cognitive function. (We also performed sub-analyses of each of these groups separately, to test whether established PD-MCI, or very early changes in the converters might be driving any changes we found). All other PD participants were classified as PD with normal cognition (PD-NC, n=50). A composite cognitive z score (averaged across the 5 individual cognitive domains) was also computed^21^.

We performed whole brain fixel based analysis using baseline age, gender and total intracranial volume as covariates, whole brain, paired comparisons of fixel-derived metrics were performed at baseline between 1) patients with PD and controls; 2) PD low versus PD high visual performers, and 3) PD-MCI versus PD-NC. Additional comparisons of interest included correlation with MoCA and two clinically-derived dementia risk scores.^22,23^ Whole brain fixel-based analysis refers to the comparison of all white matter fixels within the brain using the John Hopkins University (JHU) atlas, as is standard for this analysis.^14,24^

Additionally, to investigate longitudinal change in selective fibre pathways within the visual system, we also performed tract of interest analyses. We selected 11 white matter tracts involved in visual processing, as in our previous work^13^: Posterior thalamic and optic radiations, Splenium, Body and Genu of the corpus callosum, Superior longitudinal fasciculi, Inferior fronto-occipital fasciculi (segmentation includes the inferior longitudinal fasciculus), and Superior fronto-occipital fasciculi. We then calculated the mean FC across each tract per participant. FC was chosen for tract of interest analysis as prior works have shown it is the most sensitive fibre-specific metric of white matter degeneration in PD^13,14^. Mean FC at baseline and longitudinally was compared across PD low versus PD high visual performers. Statistical comparison was performed with a linear mixed model (using age, gender and intracranial volume as covariates). A false discovery rate (FDR) correction was performed for the 11 tracts tested, using the Benjamini/Hochberg method. Correlational analyses of mean tract FC with a combined cognitive score was performed using linear regression with age, gender and total intracranial volume as covariates (significance threshold p<0.05).

## Results

Demographics and results of clinical assessments are seen in Table 1. PD high and PD low visual performers were well matched in terms of baseline cognitive performance and MCI status, as well as disease duration, motor severity and levodopa equivalent dosage. MCI status was defined as those with MCI at baseline, and those who converted to MCI during follow-up. In total 26 patients had PD-MCI (13 at baseline and 13 converted) while the remaining 50 patients had normal cognition throughout.

**Table 1.** Demographics and results of clinical assessments.

| Characteristic | Controls<br>n= 25 | PD high<br>visual<br>performers<br>n= 54 | PD low visual<br>performers<br>n= 22 | Statistic |
| --- | --- | --- | --- | --- |
| Age (years) | **67.4 (8.2)** | **63.7 (7.5)** | **69.3 (7.6)** | **F = 7.119**<br>**p = 0.001<sup>a,b</sup>** |
| Male (%) | 12 (48) | 26 (48.1) | 15 (68.2) | $\chi^2 = 2.890$<br>$p = 0.089$ |
| Years of education | 17.8 (2.2) | 16.9 (2.9) | 18.1 (2.5) | $F = 1.919$<br>$p = 0.152$ |
| **Vision** |  |  |  |  |
| Contrast sensitivity (Pelli Robson) * | **1.8 (0.2)** | **1.8 (0.2)** | **1.7 (0.1)** | **H = 17.377**<br>**p = 0.001<sup>a,c</sup>** |
| Acuity (LogMar) * | -0.1 (0.3) | -0.1 (0.2) | -0.1 (0.1) | $H = 3.232$<br>$p = 0.199$ |
| Colour vision (D15) | 1.3 (2.4) | 2.7 (8.9) | 3.0 (4.9) | $H = 1.934$<br>$p = 0.380$ |
| **General cognition** |  |  |  |  |
| MCI | - | 13 (24.1) | 13 (59.1) | $\chi^2 = 2.933$<br>$p = 0.087$ |
| MOCA | 29.0 (1.1) | 28.3 (1.9) | 27.0 (2.7) | $H = 8.333$<br>$p = 0.016^c$ |
| MMSE | 29.2 (0.9) | 29.1 (1.7) | 28.8 (1.2) | $H = 1.534$<br>$p = 0.464$ |
| **Mood** |  |  |  |  |
| HADS anxiety | 3.5 (3.4) | 5.4 (3.7) | 6.5 (3.9) | $H = 8.201$<br>$p = 0.017^{b,c}$ |
| HADS depression | **1.2 (1.5)** | **3.2 (2.8)** | **5.7 (3.4)** | **H = 28.898**<br>**p < 0.001<sup>a,b,c</sup>** |
| **Disease specific measures** |  |  |  |  |
| Years from diagnosis | - | 3.8 (2.3) | 4.9 (2.8) | $t = -1.787$<br>$p = 0.078$ |
| UPDRS total score | - | **42.7 (16.3)** | **53.7 (28.0)** | **t = -2.102**<br>**p = 0.039** |
| UPDRS motor score | - | 22.4 (10.2) | 25.8 (16.2) | $t = -1.094$<br>$p = 0.278$ |
| LEDD | - | 401.4 (211.6) | 492.8 (225.5) | $t = -1.653$<br>$p = 0.102$ |
| RBDSQ | - | 4.0 (2.3) | 4.5 (1.9) | $t = -0.839$<br>$p = 0.404$ |

All data shown are mean (SD) except gender and MCI.
Statistical comparisons shown are across the 3 groups with subscript for post-hoc results, except for disease specific measures (comparing PD low and high visual performers). In bold characteristics that significantly differed between groups.
<sup>a</sup> Statistically significant difference between PD high visual performers and PD low visual performers, <sup>b</sup>
Statistically significant difference between PD high visual performers and controls, <sup>c</sup> Statistically significant difference between PD low visual performers and controls
\* Best binocular score used; LogMAR: lower score implies better performance, Pelli Robson: higher score implies better performance.
<sup>+</sup> Higher values imply worse image quality, <sup>-</sup> Higher values imply better image quality
MCI: mild cognitive impairment; HADS: Hospital anxiety and depression scale; MMSE: Mini-mental state examination; MOCA: Montreal cognitive assessment; UPDRS: Unified Parkinson's Disease Rating Scale, LEDD: Levodopa Equivalent Dose; RBDSQ: REM sleep behaviour disorder scale.

### Longitudinal changes in cognition

At baseline, visuospatial performance was lower, as expected, in PD low visual performers compared to PD high visual performers and controls (JLO: H=8.855, p=0.012 and Hooper: H=20.652, p<0.001). In other cognitive domains, performance at individual cognitive tasks was not significantly different between the two groups, with the exception of lower performance in PD low visual performers on a test of executive function (Stroop interference: H=6.642, p=0.036) and on a memory task (Word recognition task: H=6.716, p=0.035). Visual performance at the two computer tasks was correlated with overall cognitive performance at baseline (r=−0.306, p=0.011), as well as cognitive performance at follow up (r=−0.386, p=0.001) (Figures 2B and 2C).

**Figure 2.**
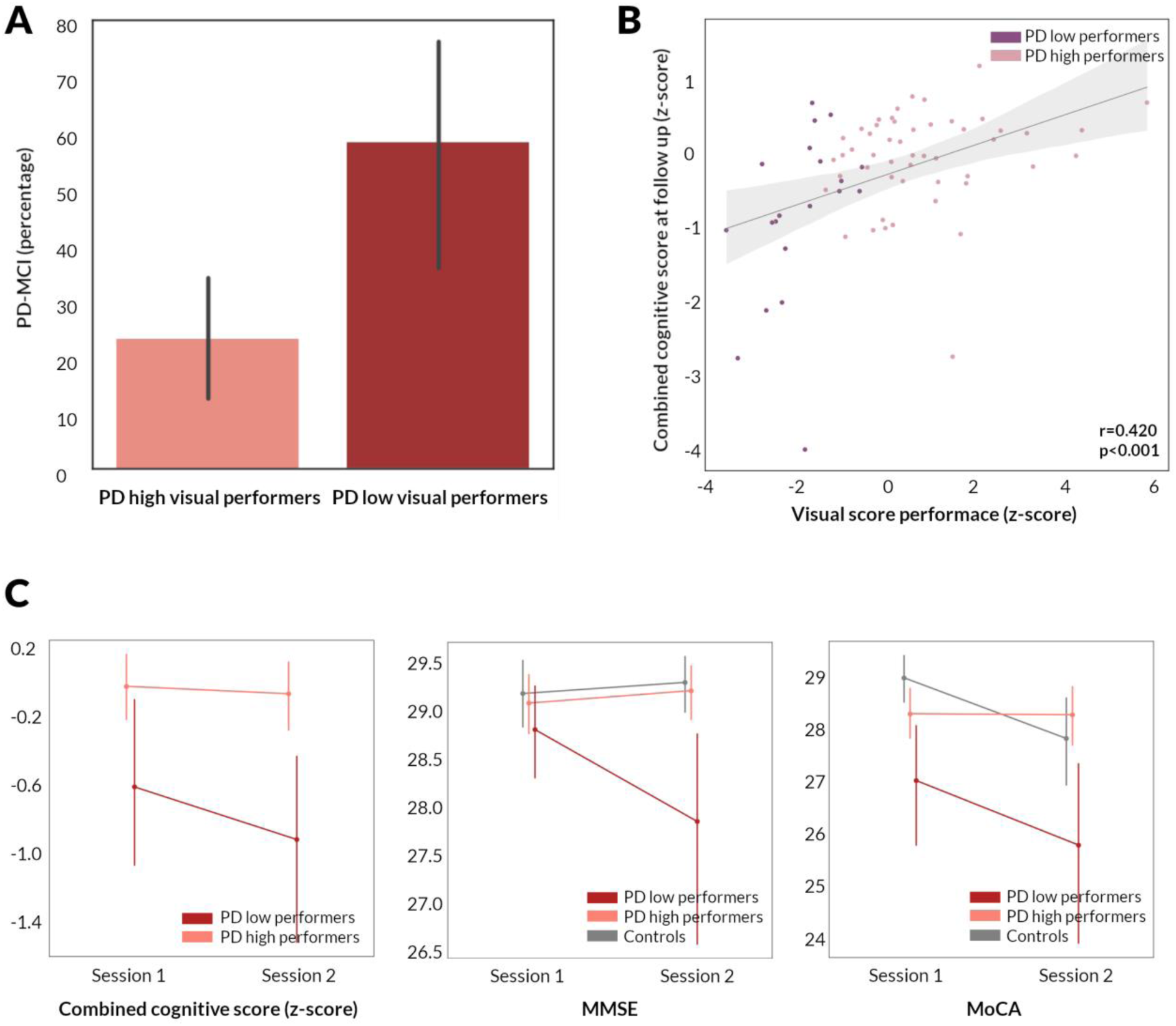
Longitudinal changes in cognition in patients with Parkinson’s disease. **A**. Percentage of patients who developed mild cognitive impairment (PD-MCI) in PD low visual performers compared to PD high visual performers. **B**. Correlation between visual performance at baseline and combined cognitive performance at 18 month follow up in patients with Parkinson’s disease (95% confidence interval). Visual performance is presented as the summed z score of the two computer-based visual tasks (Cats and Dogs task and Biological Motion task). Cognitive performance is presented as combined cognitive score (z scored against the performance of age-matched controls). **C**. Longitudinal change in combined cognitive score, MMSE and MoCA in PD low visual performers, PD high visual performers and controls. Error bars represent 95% confidence intervals. MMSE: Mini-Mental State Examination, MoCA: Montreal cognitive assessment

Significant changes in longitudinal performance between PD low and PD high visual performers were seen only for MMSE (t=−1.084, p=0.024), and Letter fluency (t=−2.825, p=0.025). However, overall cognitive performance was lower in PD low visual performers, who were more likely to develop MCI at longitudinal follow up (chi=7.031, p=0.008, Figure 2A).

### Whole-brain longitudinal changes in white matter integrity

No significant differences between patients with PD and normal controls was detected at whole brain fixel-based analysis or conventional voxel-based analysis, neither at baseline nor longitudinally.

#### 1) Baseline changes in white matter micro-structure and macro-structure related to low visual performance in Parkinson’s disease

PD low visual performers showed significant changes in white matter macro- and micro-structure compared to PD high visual performers, longitudinally (Figure 3) and already at baseline (Figure 4A), as we have shown previously ^13^. At baseline, PD low visual performers showed reduction in FC within the splenium of the corpus callosum, the right cingulum and bilateral posterior thalamic radiation. Extensive micro-structural changes (reductions in FD) were also present at baseline in PD low visual performers, with reductions within the genu, body and splenium of the corpus callosum, the right internal capsule, the cingulum bilaterally, tapetum bilaterally, posterior thalamic radiations bilaterally, right hippocampus and the right corticospinal tract. Reductions were seen across the same regions in the combined FDC metric; these were particularly pronounced within the genu and splenium of the corpus callosum with greater than 30% reductions in FDC. Figure 4A illustrates the extent of macro- and micro-structural changes seen in PD low visual performers at baseline.

**Figure 3.**
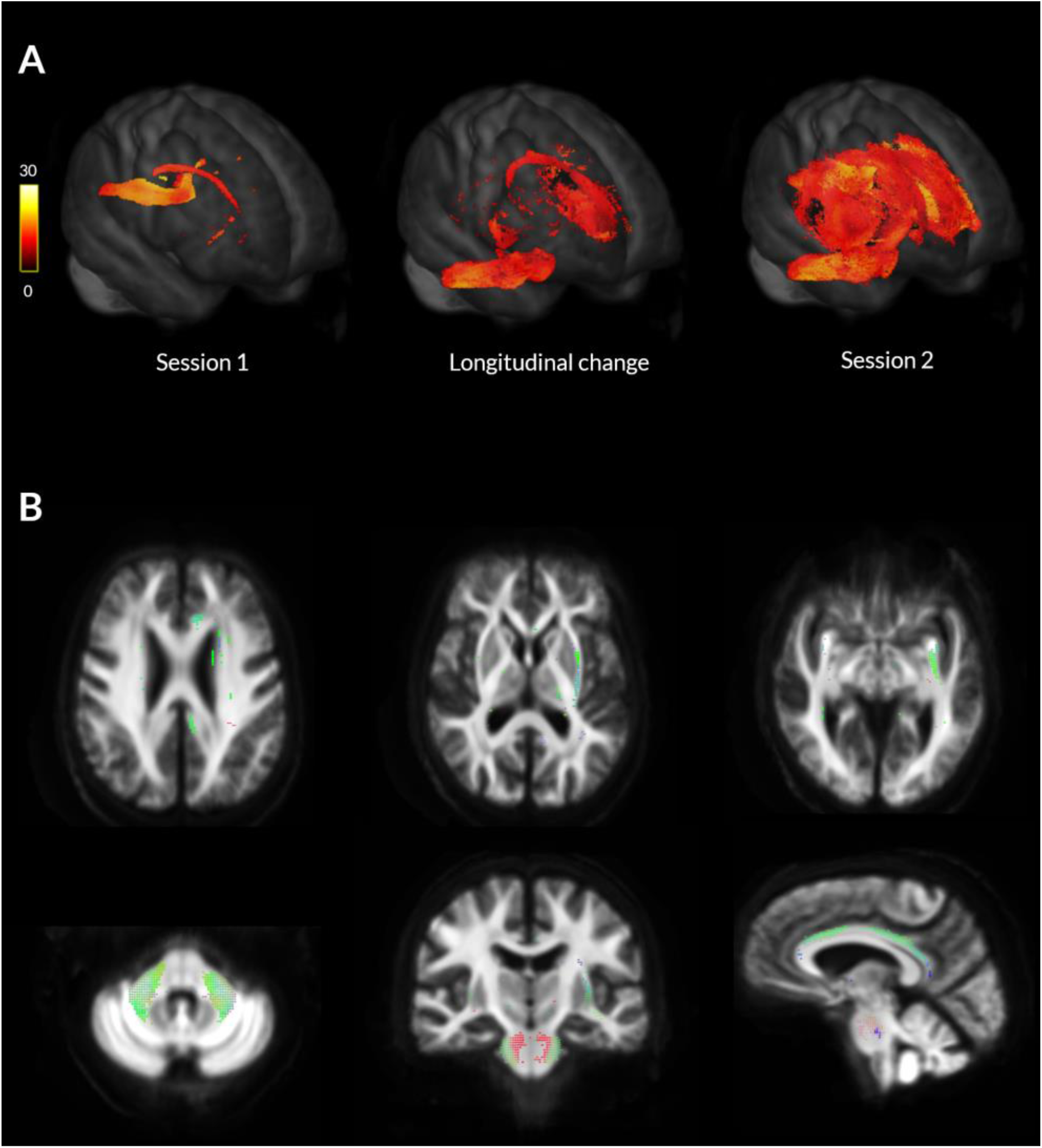
Macrostructural white matter changes in patients with Parkinson’s disease and low visual performance over time. **A**. Changes in white matter macrostructure (as seen by reduction in fibre cross-section (FC)) in PD low visual performers compared to PD high visual performers (FWE-corrected p<0.05) at Session 1 (Baseline, left), at longitudinal change (difference between the two images, middle) and at Session 2 (18 months follow up, right). Results are presented as streamlines and coloured by percentage reduction. **B**. Statistically significant (FWE-corrected p<0.05) longitudinal reductions in fibre cross-section (FC) in PD low visual performers compared to PD high visual performers. Results are coloured by direction (anterior-posterior: green; superior-inferior: blue; left-right: red).

**Figure 4.**
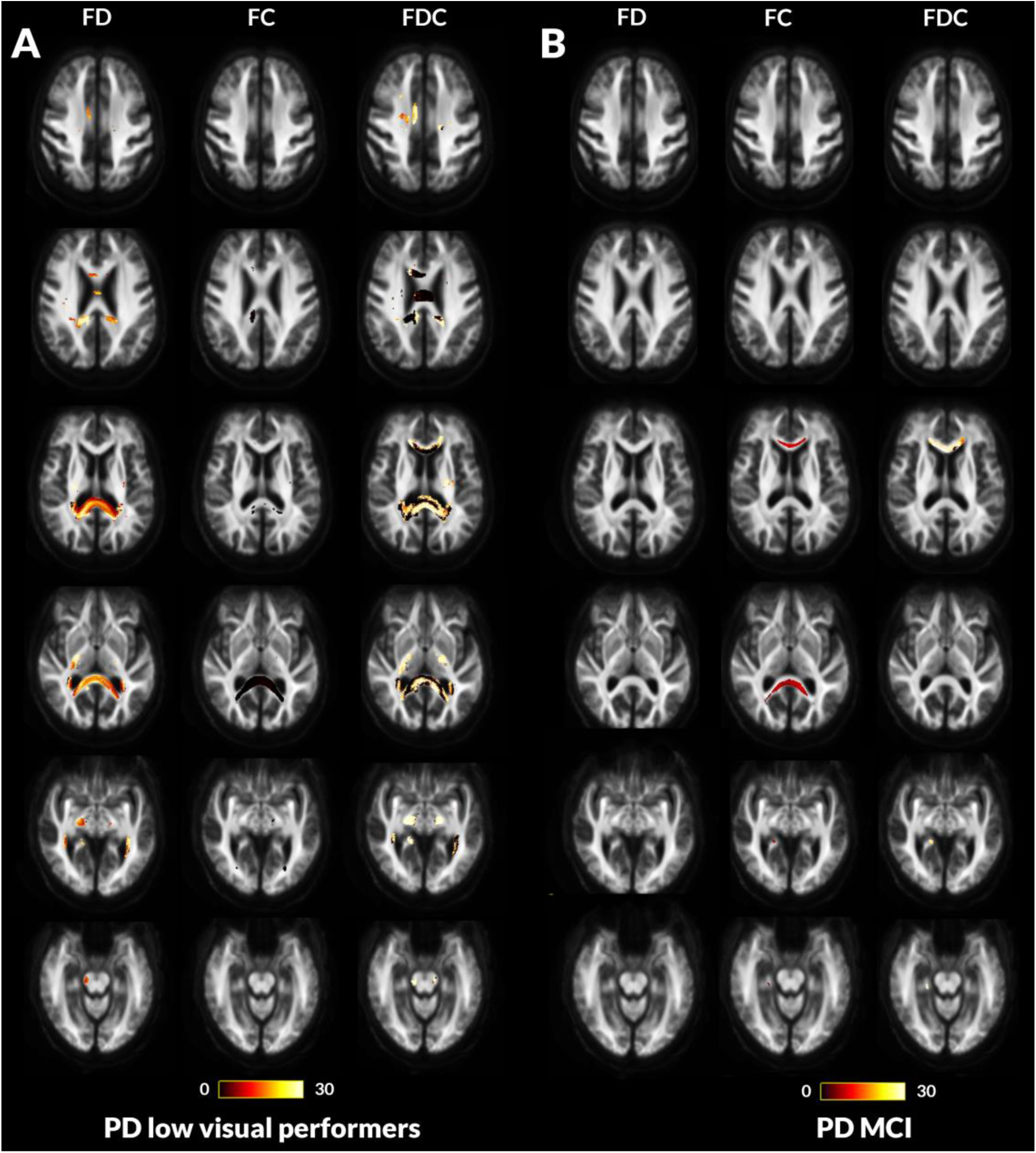
Fibre tract-specific reductions at baseline in PD low visual performers compared to PD high visual performers and PD with mild cognitive impairment compared to PD with normal cognition from whole-brain fixel-based analysis (opposite). A. PD low visual performers showed widespread microstructural (changes in fibre density (FD)) with reductions within the genu, body and splenium of the corpus callosum, the right internal capsule, the cingulum bilaterally, tapetum bilaterally, posterior thalamic radiations bilaterally, right hippocampus and the right corticospinal tract. Macrostructural changes (changes in fibre density (FC) were also seen within the splenium of the corpus callosum, the right cingulum and bilateral posterior thalamic radiations. Changes in the combined FDC metric were seen within the genu, body and splenium of the corpus callosum, the right internal capsule, the cingulum bilaterally, tapetum bilaterally, posterior thalamic radiations bilaterally, right hippocampus and the right corticospinal tract; these represent impaired overall ability to relay information in these tracts in PD low visual performers. B. Patients with Parkinson’s disease who developed mild cognitive impairment (MCI) showed macrostructural (changes in fibre cross-section (FC)) within the genu and splenium of the corpus callosum, posterior thalamic radiations bilaterally and the right hippocampus. Changes in the combined FDC metric are seen in the genu, and the right hippocampus; this represents impaired overall ability to relay information in these tracts in PD-MCI compared to PD with normal cognition (NC). No changes were seen in the FD metric for this patient group. Results are displayed as streamlines; these correspond to fixels that significantly differed between PD low and high visual performers (FWE-corrected p <0.05). Streamlines are coloured by percentage reduction (colourbars).

#### 2) White matter macro-structural changes in patients with Parkinson’s disease and mild cognitive impairment

MCI status was less sensitive than visual performance in identifying baseline white matter alterations in patients with PD, with less tracts affected in PD-MCI. PD-MCI (including with normal cognition at baseline who would convert to PD-MCI) showed reductions in FC within the genu and splenium of the corpus callosum, the right posterior thalamic radiation, as well as the right hippocampus (Figure 4B). Whilst there were no statistically significant changes in FD, the combined FDC metric showed significant, over 30%, reductions in the genu of the corpus callosum, and right hippocampus (Figure 4B). Given that white matter changes in PD-MCI and PD-low visual performers had differential spatial profiles direct comparison is difficult. However in both cases, the splenium of the corpus callosum was the most affected tract at baseline imaging. In direct comparison of mean FD, FC and FDC of the splenium, area under the curve was 0.615 for PD-MCI and 0.673 in PD-low visual performers, suggesting slightly higher sensitivity of visual performance for white matter alterations even within the same tract.

#### 3) Longitudinal white matter alterations are seen in patients with Parkinson’s disease and low visual performance

PD low visual performers showed significant changes in white matter macro-structure, with up to 22% reductions in FC compared to PD high visual performers in whole brain fixel-based analysis. These macrostructural changes were extensive and involved multiple white matter tracts, including the middle cerebellar peduncles and pontine crossing parts bilaterally, the external capsules bilaterally, the left inferior and superior fronto-occipital fasciculi, the uncinate fasciculi bilaterally, the left cingulum, the left anterior and posterior corona radiata and the genu of the corpus callosum (Figure 3), and were corrected for age as well as gender and total intracranial volume.

Importantly, cognitive measures, including MoCA scores and MCI conversion status (both assessing MCI at baseline and at follow up together and in separate sub-assessments within each group), as well as two clinical risk scores for dementia^22,23^ were not associated with longitudinal white matter changes at the micro- or at the macro-structural level.

### Voxel based analysis

Conventional voxel-based analysis of FA and MD did not show any statistically significant differences after FWE correction for PD versus controls, or PD-MCI versus PD-NC at baseline or longitudinal follow up. PD low visual performers did not show any statistically significant differences at baseline. At longitudinal imaging, PD low visual performers showed reductions in FA within the left external capsule and right posterior thalamic radiation and increase in MD within the splenium of the corpus callosum.

### Tract of interest analysis

Widespread reductions were seen in mean FC in PD low visual performers compared to PD high visual performers in tract of interest analysis. Specifically, at baseline after correction for age, gender and total intracranial volume, significant changes were seen across the corpus callosum (genu: t=−0.086, FDR-corrected q=0.002, body: t=−0.076, q=0.002, splenium: t=−0.099, q=0.002), the left inferior fronto-occipital fasciculus (t=−0.037, q=0.002) and the right superior longitudinal fasciculus (t=−0.027, q=0.002) (Figure 5B). At longitudinal follow up, all 11 selected tracts showed significant reductions (FDR-corrected) in PD low visual performers compared to PD high visual performers (Figure 5C). Across patients with PD, higher reductions in mean FC across the 11 selected tracts were associated with worsening MoCA scores (r=0.258, p=0.024) but not change in MMSE scores at 18 months follow up.

**Figure 5.**
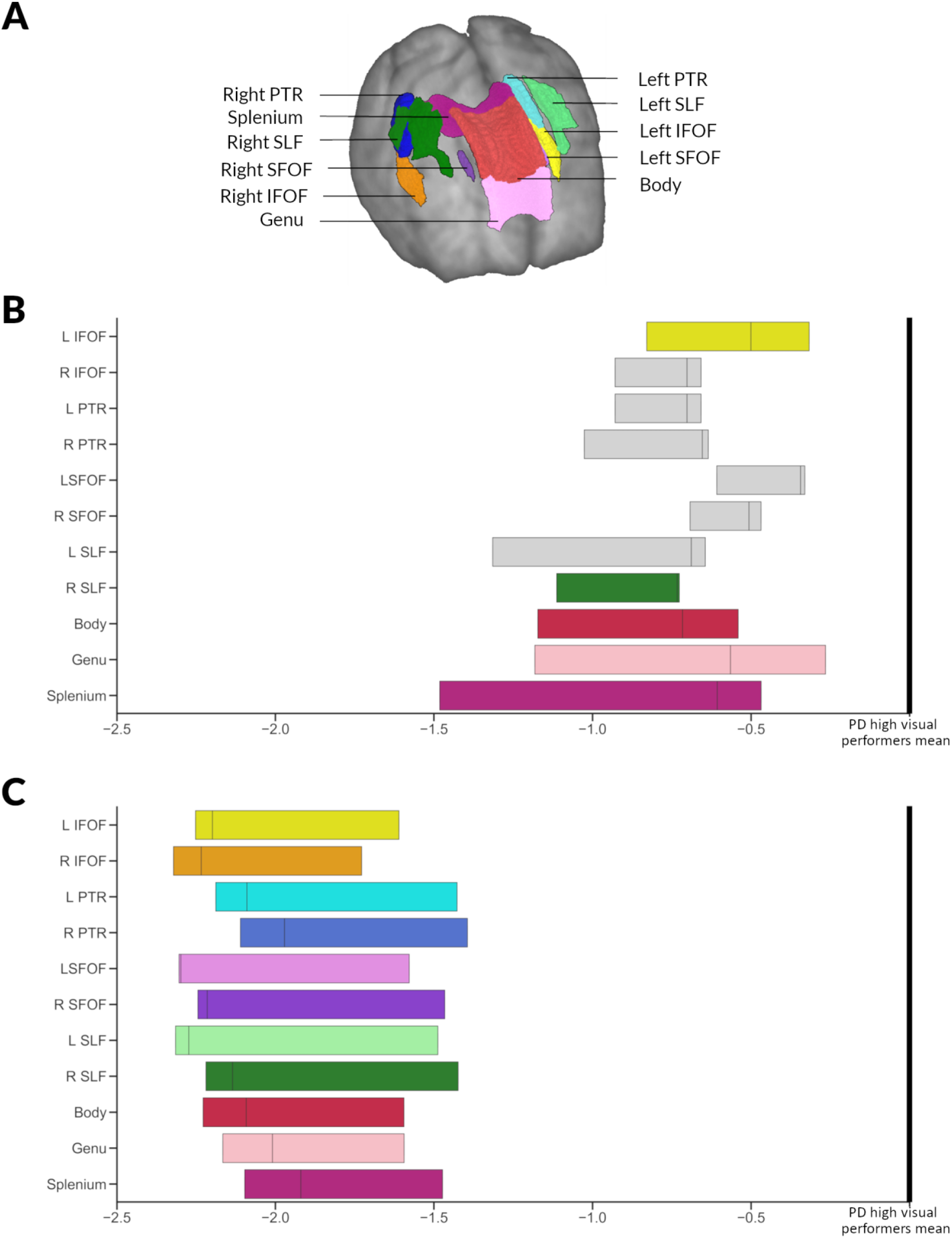
Significant tracts in Parkinson’s low performers; tract of interest analysis (Opposite). **A**. Anatomical representation of all analysed tracts. PTR: Posterior thalamic and optic radiations, SLF: Superior longitudinal fasciculi, IFOF: Inferior fronto-occipital fasciculi (segmentation includes the inferior longitudinal fasciculus), and SLF: Superior fronto-occipital fasciculi. **B**. Baseline visit. Reduction (mean, 95% CI) in fibre cross-section (FC) visualised as percentage reduction from the mean of patients with Parkinson’s disease with high visual performance. Tracts with significantly reduced FC (FDR-corrected p-value <0.05) are shown in colour whilst tracts where there are no significant changes in FDC are plotted in grey. L: Left, R: Right. **C**. Visit 2 (18 months follow up). Reduction (mean, 95% CI) in fibre cross-section (FC) visualised as percentage reduction from the mean of patients with Parkinson’s disease with high visual performance at follow-up. All 11 of the selected tracts showed significantly reduced FC (FDR corrected p-value <0.05). L: Left, R: Right.

## Discussion

Low visual performance in Parkinson’s disease was associated with cognitive decline and wide-spread white matter macrostructural changes at 18 months follow up. Specifically 1) low visual performance at baseline was correlated with worsening cognition and conversion to MCI at 18 months follow up; 2) baseline low visual performance was associated with further reductions in fibre cross-section at longitudinal follow up and 3) low performance on high-level visual tasks was associated with more extensive white matter changes at baseline than was seen in those patients who developed mild cognitive impairment on follow up. These findings provide evidence that visual changes in Parkinson’s disease are a marker for incipient white matter degeneration and cognitive decline.

### White matter alterations precede cognitive impairment in Parkinson’s disease

We have previously described the white matter changes that are associated with low visual performance at baseline^13^. We found similar but significantly less extensive white matter changes in patients who later progressed to develop MCI, specifically involving the right hippocampus and the genu and splenium of the corpus callosum. Changes within the corpus callosum, particularly its most anterior and most posterior segments, have been previously reported in PD with cognitive impairment using diffusion tensor imaging^10,25–28^. Similar changes were seen using fixel-based analysis in association with disease severity, particularly non-motor symptoms,^14^ and in patients with PD and poor visual performance^13^.

The hippocampus has a higher Lewy body count and greater cholinergic deficit in PD dementia^29^ but studies assessing grey matter atrophy and neurotransmitter changes in vivo in PD with cognitive impairment have also demonstrated changes outside the hippocampus ^30–32^, leading to debate regarding the relative importance of the hippocampus in PD dementia. In a prior meta-analysis using network-lesion mapping, our group has shown that hippocampal networks, particularly of the right hippocampus, which plays a crucial role in spatial memory^33^ are linked to PD dementia^34^. Our finding of right hippocampal tract involvement in PD-MCI provides corroborative evidence that hippocampal networks are implicated in the development of PD dementia.

In patients with low visual performance, in addition to more extensive alterations of both white matter micro-structure and morphology at baseline, additional macro-structural white matter changes developed at follow up. The right hippocampus was also affected in PD low visual performers together with its temporal lobe connections, such as the cingulum and inferior longitudinal fasciculus, which have also been implicated in Alzheimer’s disease as well as Parkinson’s with cognitive impairment^35^.

Long cortico-cortical tracts connecting frontal to occipital regions, such as the superior and inferior fronto-occipital fasciculi, and subcortical-cortical tracts, such as the external capsule, became bilaterally affected at follow up (particularly on the left hemisphere). The fronto-occipital fasciculi play a role in global cognition, attention, visual processing and executive function^36,37^. The external capsule acts as a route for cholinergic pathways and diffusion-derived metrics have been associated with cognitive performance in healthy older adults.^38^ Long-range connections play a crucial role in brain integration at network level^39^ and functional connectivity is reduced for fronto-occipital connections in PD with cognitive impairment.^40,41^

The cerebellar tracts, particularly the middle cerebellar peduncles showed significant macro-structural changes at follow up in PD low visual performers. The cortico-ponto-cerebellar pathway, which connects cortical regions to the cerebellum via the contralateral middle cerebellar peduncle, may play a key role in cognition^42,43^. In patients with Parkinson’s disease, functional connectivity between the vermis and visual association and pre-frontal cortex is weaker than controls and correlates with cognitive performance, even in the absence of cerebellar volume loss or changes in cortical thickness^21^. Although traditionally regarded as unaffected by alpha-synuclein pathology and excluded from the Braak Lewy body pathological staging, recent evidence suggests that cerebellar nuclei and surrounding white matter do accumulate alpha-synuclein aggregates^44^. Our findings provide further support to structural white matter changes within the cerebellum in patients with PD and cognitive impairment.

In patients with PD and poor visual performance, we saw widespread white matter alterations, with interhemispheric connections in the corpus callosum, long fronto-occipital and hippocampal connections affected first, followed by diffuse macro-structural changes across multiple association tracts. These diffuse changes may explain the alterations in global functional and structural connectivity seen in Parkinson’s disease^45–47^ and emphasise the new insights that network approaches can provide in our understanding of Parkinson’s dementia.

### Impending cognitive decline and macrostructural white matter degeneration in patients with Parkinson’s disease and visual dysfunction: an opportunity for intervention

Baseline visual performance was associated with cognitive performance at follow-up in patients with PD; and poor visual performance predicted worsening cognition and the development of MCI. Prior work from our group has shown that visuo-perceptual deficits are linked to poorer cognitive performance at baseline and clinical scores for dementia^5,16^ as well as worsening MoCA at 1 year follow up^17^. This imaging study provides further evidence of impending cognitive impairment in patients with PD and visuo-perceptual deficits.

We also found that PD low visual performers exhibited significant and widespread reduction in fibre cross section, signalling widespread macro-structural white matter alterations at 18 months follow up. In contrast, patients with PD who are classified as high dementia risk by two clinically-derived scores^22,22^ as well as those who developed MCI at follow up did not exhibit further white matter alterations, with PD-MCI showing established white matter degeneration at baseline. These findings raise the possibility that poor visual performance could represent a window of opportunity for therapeutic interventions in patients with PD, by identifying patients with imminent but not yet clinically apparent cognitive impairment and white matter degeneration.

### Limitations and future directions

Several considerations need to be taken into account when interpreting the results from our study. All raw imaging data in our cohort were visually inspected and clinically significant cerebrovascular disease was excluded, however, due to the acquired MRI sequences, we could not systematically quantify and control for white matter hyperintensities, which could lead to fibre density reduction^48^. This is in line with other published studies using fixel-based analysis,^14,24,49,50^ but future studies need to clarify the effect of white matter hyperintensities on fixel-based metrics. At both study timepoints, participants with PD underwent imaging on their usual dopaminergic medications. Corrected fractional anisotropy is not affected by levodopa^51^ therefore it is unlikely that dopaminergic medication would affect fixel-based metrics. In addition, levodopa equivalent doses were not significantly different between PD low and PD high visual performers at any study time-point. Follow up imaging and psychological testing was performed at 18 months; longer follow up times of patients with PD who will proceed to develop dementia could provide further insights to the temporal order of white matter degeneration in PD with cognitive impairment.

## Conclusions

Poor visual function in Parkinson’s disease is associated with worsening cognition and conversion to MCI at 18 months follow up. Poor visual performance was also associated with diffuse white matter macro-structural changes at follow up. These findings provide insights into the temporal pattern of white matter degeneration associated with cognitive impairment in Parkinson’s disease and highlight an at-risk population for possible therapeutic intervention.

## Acknowledgements

We gratefully acknowledge the support of NVIDIA Corporation with the donation of the Quadro P6000 GPU used for this research. The authors acknowledge the use of the UCL Myriad High Performance Computing Facility (Myriad@UCL), and associated support services, in the completion of this work. This research was also supported by the National Institute for Health Research University College London Hospitals Biomedical Research Centre.

## Author Contributions

Study design and concept: AZ, RW, data collection: AZ, LAL, RW, imaging and statistical analysis: AZ, drafting and revision of the manuscript: AZ, PMC, LAL, AL, RW.

